# Contribution of markers of adiposopathy and adipose cell size in predicting insulin resistance in women of varying age and adiposity

**DOI:** 10.1101/2021.11.03.467138

**Authors:** Eve-Julie Tremblay, André Tchernof, Mélissa Pelletier, Nicolas Chabot, Denis R. Joanisse, Pascale Mauriège

## Abstract

Adipose tissue (AT) dysfunctions, such as adipocyte hypertrophy, macrophage infiltration and secretory adiposopathy (low plasma adiponectin/leptin, A/L, ratio), associate with metabolic disorders. However, no study has compared the relative contribution of these markers to cardiometabolic risk in women of varying age and adiposity. Body composition, regional AT distribution, lipid-lipoprotein profile, glucose homeostasis and plasma A and L levels were determined in 67 women (age: 40-62 years; BMI: 17-41 kg/m^2^). Expression of macrophage infiltration marker CD68 and adipocyte size were measured from subcutaneous abdominal (SCABD) and omental (OME) fat. AT dysfunction markers correlated with most lipid-lipoprotein levels. The A/L ratio was negatively associated with fasting insulinemia and HOMA-IR, while SCABD or OME adipocyte size and SCABD CD68 expression were positively related to these variables. Combination of tertiles of largest adipocyte size and lowest A/L ratio showed the highest HOMA-IR. Multiple regression analyses including these markers and TAG levels revealed that the A/L ratio was the only predictor of fasting insulinemia and HOMA-IR. The contribution of the A/L ratio was superseded by adipose cell size in the model where the latter replaced TAGs. Finally, leptinemia was a better predictor of IR than adipocyte size and the A/L ratio in our participant sample.

## Introduction

Obesity contributes to the development of several metabolic complications, although the distribution of adipose tissue (AT) is more significant with respect to health outcomes than obesity per se [1,2]. Indeed, visceral obesity, defined as an accumulation of intra-abdominal AT, is closely associated with metabolic disorders such as chronic, low-grade inflammation, insulin resistance (IR), metabolic syndrome (MetS), type 2 diabetes (T2D), dyslipidemia, hypertension and cardiovascular disease (CVD), *i.e*. increased cardiometabolic risk [3,4].

In addition, the way in which AT manages the excess energy as lipids can vary with obesity and leads to greater health risk [5,6]. In this regard, AT expansion can occur via either adipocyte hypertrophy or hyperplasia [7], and the number and the size of adipose cells may represent significant markers of AT metabolic dysfunction [8,9]. While cell size is known to generally increase with obesity level irrespective of the anatomical location [10], hyperplasia was predominant in subcutaneous abdominal (SCABD) fat depots, whereas adipose cell hypertrophy was observed both in the omental (OME) and SCABD compartments of women with obesity [11]. Moreover, women characterized by OME adipocyte hypertrophy presented a worsened lipid-lipoprotein profile compared with women characterized by OME adipocyte hyperplasia, regardless of differences in body composition and regional fat distribution [9]. These results suggest that adipose cell size, and more importantly OME adipocyte hypertrophy, more closely associates with cardiometabolic disorders.

With obesity, AT is also infiltrated by immune cells such as macrophages which contribute to local and systemic inflammation [12], and are thus involved in metabolic complications such as IR [13]. This infiltration is typically assessed by the expression of several macrophage-specific genes. More specifically, M1 pro-inflammatory macrophages express the integrin α-chain CD11c, as well as CD11b, while M2 anti-inflammatory macrophages express only CD11b [14]. CD68 (a marker of total macrophage inflammation), CD11b, and CD11c are surface markers involved in the binding of antigens, adhesion molecules, and macrophage-specific growth factors [15]. Moreover, the CD68 transcript is a well-established marker of total macrophage infiltration, and mean CD68+ cell percentage is considered as the proportion of resident cells at the junction of two or more adipocytes in SCABD and/or OME AT [16,17]. Total adiposity, regional fat distribution, OME and SCABD adipose cell sizes are all positively correlated with the expression of CD68, CD11b, and CD11c, stronger correlations being observed at the SCABD than the OME level [16,17].

Another dimension of adiposopathy (or “sick fat”) relates to secretory dysfunctions [18], which are frequently observed with obesity. Indeed, AT secretes a wide variety of biologically active adipokines [19], including leptin and adiponectin which are primarily secreted by subcutaneous and visceral adipose cells respectively [20,21] To clearly distinguish this dimension of adiposopathy from other AT dysfunctions, we have chosen to name it secretory adiposopathy. The adiponectin (anti-inflammatory) to leptin (pro-inflammatory) (A/L) ratio is often used to assess this dimension [22]. Indeed, as an increase in circulating leptin and a decrease in adiponectin levels are observed in obesity [23], a low plasma A/L ratio is associated with a worsened AT secretory dysfunction [24]. Also, the A/L ratio was shown to be a stronger correlate to IR than adiponectin or leptin alone [25], an independent predictor of insulin sensitivity (IS) in women with moderate obesity [26] and men presenting obesity without [24] or with MetS [27], glucose intolerance [28] or T2D [25].

To the best of our knowledge, there has been no attempt to compare the relative importance of the secretory adiposopathy to other AT dysfunction markers regarding cardiometabolic risk. Therefore, this study aimed to examine the respective contribution of the plasma A/L ratio, adipose cell size and the expression of AT macrophage infiltration markers at both the SCABD and OME levels on adiposity, lipid-lipoprotein profile, glucose homeostasis, and more particularly IR, in a sample of women with varying age and adiposity.

## Results

### Participants’ characteristics

**Table 1** summarizes participants’ characteristics. Although being overweight on average according to their BMI (27.2 ± 5.0 kg/m^2^) (means ± SD), patients nevertheless showed a wide range of body fatness, regional fat distribution and adipocyte hypertrophy at both the SCABD and the OME levels. From the range of fasting glucose and insulin values, our sample included normal glucose tolerant to intolerant (but not diabetic) individuals. Also, adiponectin and leptin levels showed marked inter-individual variation attesting to variations in AT secretory function, with plasma A/L ratio values ranging from 0.04 to 34.2.

**Table 1.** Characteristics of the women studied.

|  | n | Mean (SD) | Range (min - max) |
| --- | --- | --- | --- |
| Age (years) | 67 | 47 (5) | 40 - 62 |
| *Anthropometry and body fatness* |  |  |  |
| Body weight (kg) | 65 | 70.6 (14.8) | 48.5 - 110.5 |
| Body mass index (kg/m<sup>2</sup>) | 65 | 27.2 (5.0) | 17.2 - 41.3 |
| Body fat mass (kg) | 65 | 25.5 (9.0) | 10.0 - 50.8 |
| Body fat percentage (%) | 65 | 35.1 (6.0) | 32.5 - 63.0 |
| Lean body mass (kg) | 65 | 43.1 (6.6) | 19.6 - 47.5 |
| *Abdominal adipose tissue areas (cm<sup>2</sup>)* |  |  |  |
| Total | 63 | 424 (180) | 128 - 991 |
| Subcutaneous | 63 | 328 (141) | 94 - 759 |
| Visceral | 63 | 97 (46) | 34 - 233 |
| *Adipose cell size (μm)* |  |  |  |
| Subcutaneous abdominal | 63 | 98.5 (12.9) | 66.8 - 122.7 |
| Omental | 59 | 80.8 (16.2) | 51.7 - 118.7 |
| *Lipid-lipoprotein profile* |  |  |  |
| Cholesterol (mmol/L) | 62 | 4.81 (0.64) | 3.43 - 6.12 |
| LDL-cholesterol (mmol/L) | 61 | 2.71 (0.59) | 1.31 - 3.96 |
| HDL-cholesterol (mmol/L) | 62 | 1.49 (0.39) | 0.81 - 2.69 |
| Cholesterol/HDL-C | 62 | 3.43 (0.93) | 1.64 - 6.22 |
| Triacylglycerols (mmol/L) | 62 | 1.34 (0.74) | 0.51 - 4.69 |
| *Glucose homeostasis* |  |  |  |
| Fasting glucose (mmol/L) | 63 | 5.6 (0.6) | 4.6 - 7.8 |
| Fasting insulin (pmol/L) | 63 | 11.4 (5.6) | 3.4 - 27.6 |
| HOMA-IR | 63 | 2.9 (1.7) | 0.8 - 8.8 |
| *Adipokines* |  |  |  |
| Adiponectin (μg/mL) | 63 | 10.8 (5.8) | 0.7 - 28.6 |
| Leptin (ng/mL) | 61 | 26.4 (20.3) | 0.5 - 72.4 |
| Adiponectin/Leptin (10<sup>-3</sup>) | 60 | 2.7 (6.9) | 0.04 - 34.2 |
| *Macrophage infiltration markers* |  |  |  |
| Subcutaneous abdominal |  |  |  |
| CD68 mRNA | 51 | 1.26 (0.77) | 0.39 - 5.03 |
| CD11b mRNA | 52 | 0.96 (0.54) | 0.25 - 2.53 |
| CD11c mRNA | 52 | 1.72 (1.39) | 0.22 - 7.08 |
| Omental |  |  |  |
| CD68 mRNA | 53 | 1.17 (0.53) | 0.46 - 3.13 |
| CD11b mRNA | 53 | 0.98 (0.54) | 0.30 - 2.54 |
| CD11c mRNA | 54 | 0.84 (0.68) *** | 0.19 - 2.92 |

SD, standard deviation. C, cholesterol; HDL, high-density lipoprotein; CD: cluster designation; LDL, low-density lipoprotein; HOMA-IR, HOMeostatic Model Assessment of Insulin Resistance.
\*\*\*compared to subcutaneous abdominal at p<0.0005.

Regarding AT macrophage infiltration markers, CD68 mRNA levels were similar in SCABD and OME fat. Similarly, expression of CD11b did not show regional variation, although CD11c mRNA abundance was higher in SCABD than in OME AT.

### Relationships between selected AT dysfunction markers and participants’ characteristics

The plasma A/L ratio was strongly and negatively associated with both the SCABD and VISC AT areas (−0.80≤rho≤-0.70; p≤0.0005). Positive correlations were observed between adipose cell sizes and abdominal AT areas (0.70≤rho≤0.78; p≤0.0005). At the SCABD fat level, CD68 expression was positively associated with VISC AT area while CD11b mRNA abundance was positively associated with all AT areas. At the OME level, CD11b and CD11c mRNA levels were positively related to VISC fat area (data not shown).

As depicted in **Table 2**, the A/L ratio was also positively associated with HDL-cholesterol and negatively with TAG levels, but not with total cholesterol or LDL-cholesterol concentrations. Both SCABD and OME adipose cell sizes were negatively associated with HDL-cholesterol and positively with the cholesterol/HDL-cholesterol ratio as well as with TAGs. OME adipocyte size showed a marginal positive correlation with LDL-cholesterol. SCABD CD68 expression was positively associated with most lipid-lipoprotein levels, except cholesterol and HDL-cholesterol, whereas OME CD68 was negatively correlated with HDL-cholesterol levels and positively correlated with the cholesterol/HDL-cholesterol ratio. In addition, SCABD CD11b mRNA levels were positively associated with the cholesterol/HDL-cholesterol ratio, while being negatively related to HDL-cholesterol concentrations. Positive associations were found between OME CD11b expression and the cholesterol/HDL-cholesterol ratio. Finally, there was no relationship between CD11c expression in each fat depot and the lipid-lipoprotein profile.

**Table 2.** Spearman correlations between several AT dysfunction markers and patients’ lipid-lipoprotein profile.

|  | Plasma<br>A/L ratio<br>(n = 55-58) | SCABD<br>adipose<br>cell size<br>(n = 56-57) | OME<br>adipose<br>cell size<br>(n = 54-56) | SCABD<br>CD68<br>mRNA<br>(n = 47) | OME<br>CD68<br>mRNA<br>(n = 49) | SCABD<br>CD11b<br>mRNA<br>(n = 48) | OME<br>CD11b<br>mRNA<br>(n = 49-50) | SCABD<br>CD11c<br>mRNA<br>(n = 48) | OME<br>CD11c<br>mRNA<br>(n = 50) |
| --- | --- | --- | --- | --- | --- | --- | --- | --- | --- |
| Cholesterol (mmol/L) | -0.13 | 0.13 | 0.16 | 0.24 | -0.06 | 0.08 | 0.17 | 0.15 | 0.10 |
| LDL-cholesterol (mmol/L) | -0.16 | 0.22 | 0.28* | 0.32* | 0.14 | 0.13 | 0.23 | 0.20 | 0.15 |
| HDL-cholesterol (mmol/L) | 0.38** | -0.48*** | -0.48*** | -0.28*** | -0.37* | -0.34* | -0.27 | -0.25 | -0.14 |
| Cholesterol/HDL-C | -0.43** | 0.52*** | 0.54*** | 0.31*** | 0.35* | 0.33* | 0.34* | 0.27 | 0.23 |
| Triacylglycerols (mmol/L) | -0.46*** | 0.46*** | 0.37** | 0.30* | 0.16 | 0.26 | 0.26 | 0.11 | 0.09 |

SCABD: subcutaneous abdominal; OME: omental. For other abbreviations, see legends to Table 1. \*\*\*p<0.0005, \*\*p<0.005, \*p<0.05.

Regarding glucose homeostasis, adipose cell size, irrespective of anatomic location, was positively related to fasting insulin levels and HOMA-IR **(Figures 1A to 1D)** while the plasma A/L ratio was negatively associated with the latter variables **(Figures 1E and 1F).** The apparently stronger association between A/L ratio and SCABD adipocyte size in predicting IR led us to investigate the relationships between secretory adiposopathy and AT morphology. We ran a partial correlation analysis between the A/L ratio and SCABD or OME adipose cell size. In these analyses, the correlation between A/L ratio and SCABD adipose cell size was retained (partial r=-0.50; p<0.0005), and that with OME cell size was lost (partial r=-0.15; p=0.36).

**Figure 1.**
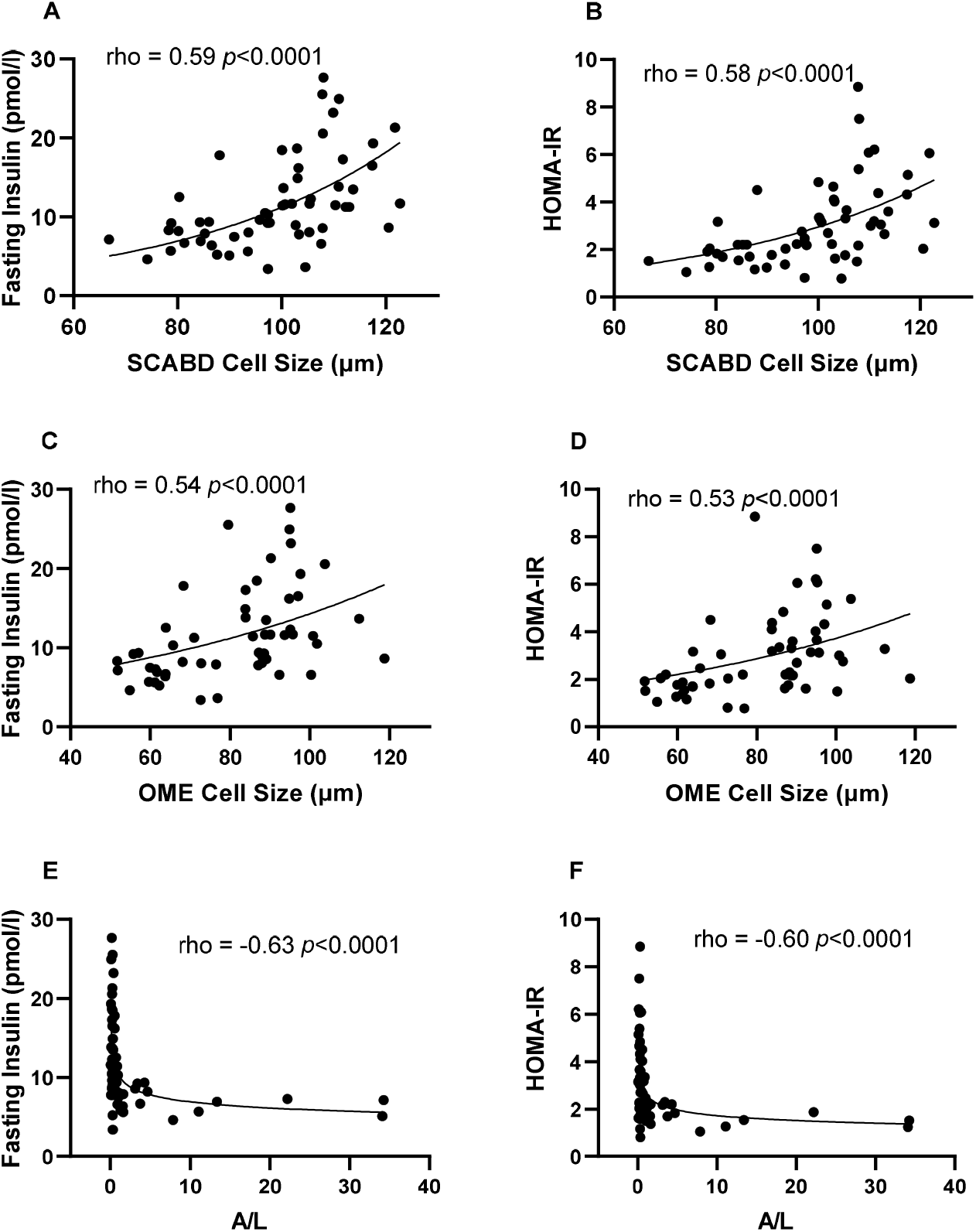
Relationships between subcutaneous abdominal (panels A and B) or omental (panels C and D) adipose cell size, plasma A/L ratio (panels E and F) and fasting insulin levels (left) and HOMA-IR index (right). A: adiponectin; HOMA-IR: HOmeostasic Model Assessment of Insulin Resistance; L: leptin; OME: omental; SCABD: subcutaneous abdominal.

Regarding AT macrophage infiltration, SCABD CD68 mRNA abundance was positively associated with fasting insulin levels and HOMA-IR **(Figures 2A and 2B),** in contrast to OME CD68 mRNA abundance **(Figures 2C and 2D). In addition,** SCABD CD11b expression was positively related to fasting insulinemia and HOMA-IR **(Figures 2E and 2F)**, but not that of OME **(Figures 2G and 2H)**. CD11c expression assessed either at the SCABD or the OME level was not associated with any variable reflecting glucose homeostasis (data not shown).

**Figure 2.**
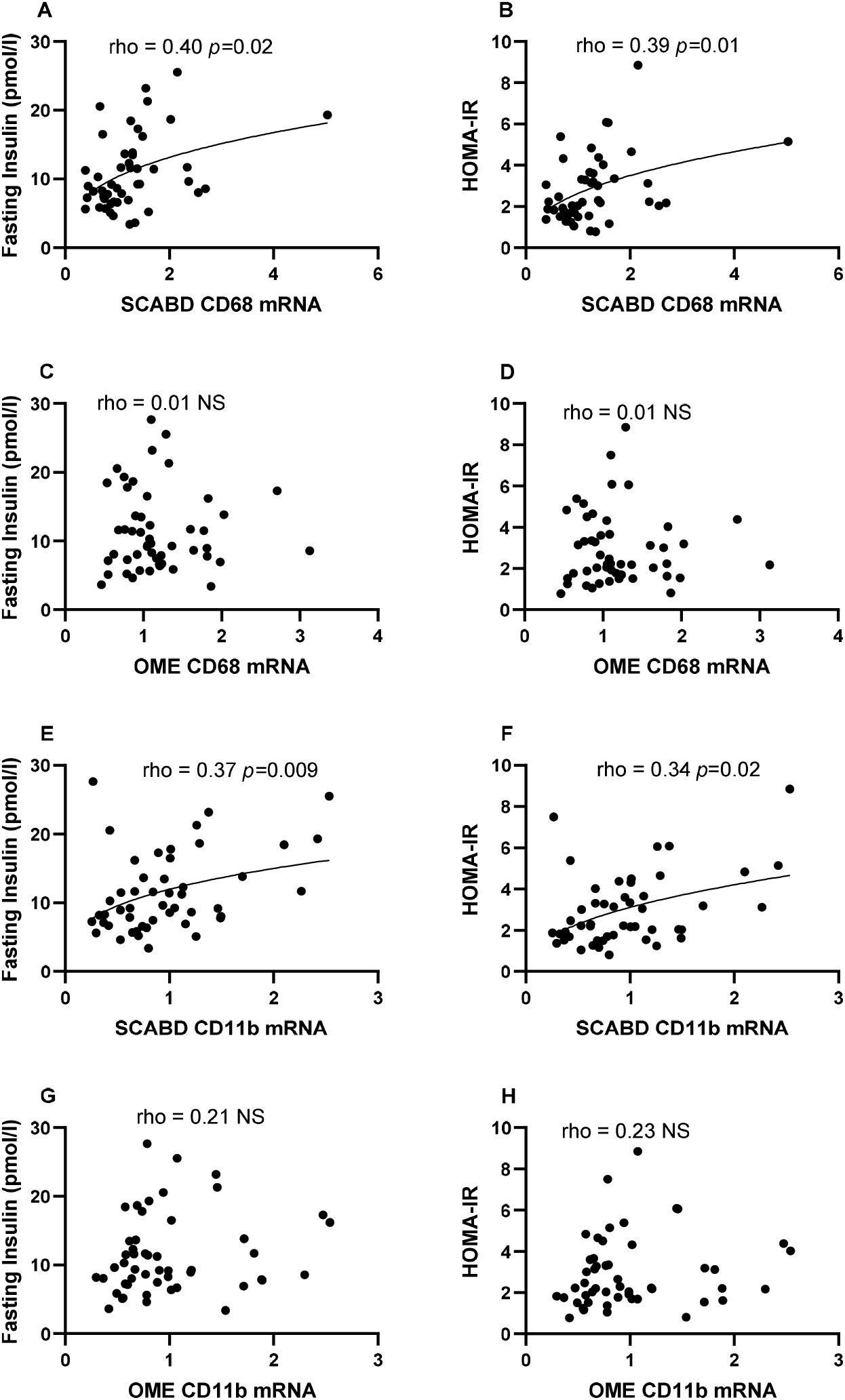
Relationships between subcutaneous abdominal CD68 (panels A and B) and CD11b (panels E and F) mRNA levels or omental CD68 (panels C and D) and CD11b (panels G and H) mRNA levels and fasting insulin levels (left) and HOMA-IR (right). For abbreviations, see legends to Figure 1 and Table 1.

Finally, fasting glucose levels were related to SCABD or OME adipose cell size (0.38<rho< 0.43; 0.001<p<0.005) but were associated neither with the plasma A/L ratio (rho =-0.24; NS) nor to CD68 expression, irrespective of the fat depot (0.02<rho<0.24; NS).

To further investigate the relationships between AT morphology and secretory adiposopathy, we examined the distribution of adipose cell size in low (0.4 ± 0.8) vs high (5.1 ± 10.1) (means ± SD) tertiles of plasma A/L ratio **(Figure 3)**. As shown in the inset Table, participants with secretory adiposopathy (*i.e*., a low A/L ratio) were characterized as having higher BMI and % fat, greater subcutaneous abdominal and visceral AT areas as well as larger SCABD **(Figure 3A)** and OME **(Figure 3B)** adipocytes (0.0005<p<0.0001), compared to those with a high A/L ratio.

**Figure 3.**
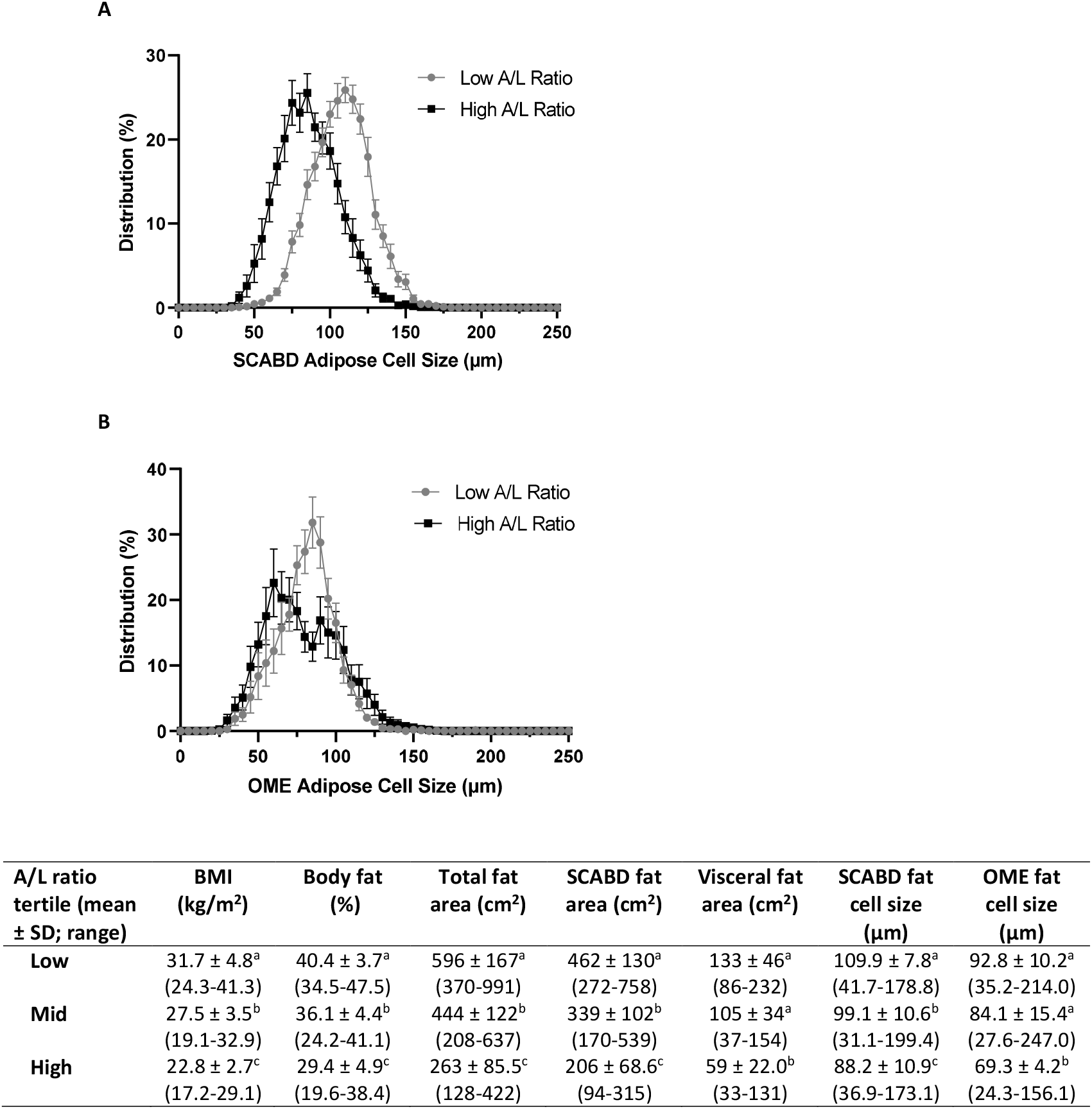
Subcutaneous abdominal (panel A) and omental (panel B) adipose cell size distributions in low vs. high plasma A/L ratio tertiles. The inset Table shows body composition, regional fat distribution and adipose cell size in the low, mid, and high A/L ratio tertiles. Values not sharing a same letter are statistically different from each other at p values ranging from 0.0001 and 0.01. For abbreviations see legends to Figure 1.

Despite these differences, we observed a large overlap in the distribution curves of cell sizes of those with high or low secretory adiposopathy. Thus, we further examined, using 3-D graphs of tertiles, whether the combination of low plasma A/L ratio and adipocyte hypertrophy could be related to increased HOMA-IR.Indeed, the HOMA-IR index was increased by 2-3-fold compared to its value when considering the combination of a low A/L ratio and a high SCABD (**Figure 4A**) or OME (**Figure 4B**) adipose cell size. These data show that a combination of these two factors associates with the highest level of IR in our sample of women.

**Figure 4.**
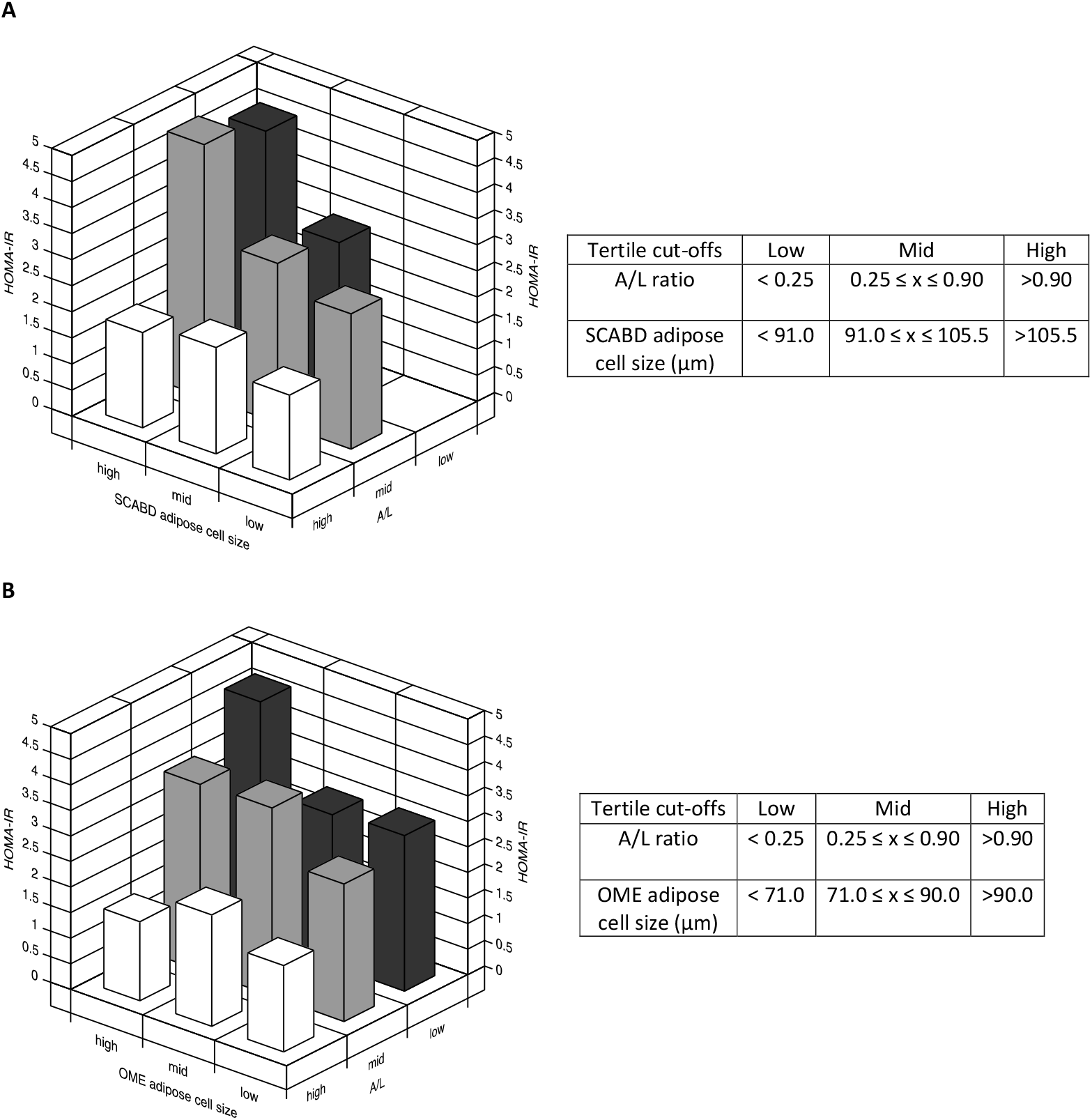
Combination of adiposopathy (plasma A/L ratio) and subcutaneous abdominal (panel A) or omental (panel B) adipose cell size tertiles in relation to the HOMA-IR index. Values of tertiles are shown in the corresponding tables. For abbreviations see legends to Figure 1.

On the other hand, a strong and negative relationship was observed between the A/L ratio and SCABD or OME adipose cell size (−0.72<rho<-0.62; p<0.0001). As these relationships did not reach collinearity, we proceeded with multiple regression analyses to identify variables independently predicting IR in our sample of women.

### Multiple regression modelling of glucose homeostasis

To investigate among selected variables the best predictors of glucose homeostasis and more particularly of IR, the strongest single correlates from different dimensions of AT dysfunction were entered into stepwise multiple regression models with or without SCABD or OME adipose cell size **(Tables 3)**. In models without cell size, the plasma A/L ratio was a strong predictor of HOMA-IR and fasting insulin levels but not of fasting glucose concentrations, TAGs being the best predictor of the latter **(Table 3A)**. When TAGs were replaced by adipose cell size, the A/L ratio lost its independent predicting ability of the different variables reflecting glucose homeostasis **(Table 3B)**. SCABD or OME adipocyte size was the only predictor of HOMA-IR, fasting insulin and glucose levels retained in these models.

**Tables 3.** Multiple regression models of predictors of glucose homeostasis.

**A. Model including AT dysfunction markers without adipose cell size**
| SCABD |  |  |  | OME |  |  |  |
| --- | --- | --- | --- | --- | --- | --- | --- |
| HOMA index (IR) |  |  |  |  |  |  |  |
| | Model $R^2 = 0.38$ | | | | Model $R^2 = 0.40$ | | |
| | $\beta$ | Partial $R^2$ | $p$ | | $\beta$ | Partial $R^2$ | $p$ |
| A/L ratio | -0.428 | 0.32 | 0.0055 | A/L ratio | -0.445 | 0.31 | 0.0021 |
| Triacylglycerols | 0.180 | 0.04 | NS | Triacylglycerols | 0.315 | 0.05 | 0.0277 |
| CD68 | 0.168 | 0.02 | NS | CD68 | -0.185 | 0.04 | NS |

| Fasting insulin |  |  |  |  |  |  |  |
| --- | --- | --- | --- | --- | --- | --- | --- |
| | Model $R^2 = 0.36$ | | | | Model $R^2 = 0.15$ | | |
| | $\beta$ | Partial $R^2$ | $p$ | | $\beta$ | Partial $R^2$ | $p$ |
| A/L ratio | -0.492 | 0.33 | 0.0005 | A/L ratio | -0.475 | 0.32 | 0.0011 |
| CD68 | 0.217 | 0.03 | NS | Triacylglycerols | 0.282 | 0.05 | 0.0467 |
| Triacylglycerols | NE | - | - | CD68 | -0.187 | 0.03 | NS |

| Fasting glucose |  |  |  |  |  |  |  |
| --- | --- | --- | --- | --- | --- | --- | --- |
| | Model $R^2 = 0.14$ | | | | Model $R^2 = 0.15$ | | |
| | $\beta$ | Partial $R^2$ | $p$ | | $\beta$ | Partial $R^2$ | $p$ |
| Triacylglycerols | 0.374 | 0.14 | 0.0033 | Triacylglycerols | 0.374 | 0.15 | 0.0033 |
| A/L ratio | NE | - | - | A/L ratio | NE | - | - |
| CD68 | NE | - | - | CD68 | NE | - | - |

**B. Model including AT dysfunction markers with adipose cell size**
| SCABD |  |  |  | OME |  |  |  |
| --- | --- | --- | --- | --- | --- | --- | --- |
| HOMA index (IR) |  |  |  |  |  |  |  |
| | Model $R^2 = 0.45$ | | | | Model $R^2 = 0.36$ | | |
| | $\beta$ | Partial $R^2$ | $p$ | | $\beta$ | Partial $R^2$ | $p$ |
| Adipose cell size | 0.552 | 0.43 | <0.0001 | Adipose cell size | 0.391 | 0.30 | 0.0326 |
| CD68 | 0.139 | 0.02 | NS | A/L ratio | -0.266 | 0.03 | NS |
| A/L ratio | NE | - | - | CD68 | -0.154 | 0.03 | NS |

**Fasting insulin**
| | Model $R^2 = 0.44$ | | | | Model $R^2 = 0.37$ | | |
| --- | --- | --- | --- | --- | --- | --- | --- |
| | $\beta$ | Partial $R^2$ | $p$ | | $\beta$ | Partial $R^2$ | $p$ |
| Adipose cell size | 0.540 | 0.42 | 0.0002 | Adipose cell size | 0.397 | 0.31 | 0.0288 |
| CD68 | 0.130 | 0.02 | NS | A/L ratio | -0.272 | 0.04 | NS |
| A/L ratio | NE | - | - | CD68 | -0.157 | 0.02 | NS |

**Fasting glucose**
| | Model $R^2 = 0.23$ | | | | Model $R^2 = 0.09$ | | |
| --- | --- | --- | --- | --- | --- | --- | --- |
| | $\beta$ | Partial $R^2$ | $p$ | | $\beta$ | Partial $R^2$ | $p$ |
| Adipose cell size | 0.414 | 0.23 | 0.0014 | Adipose cell size | 0.316 | 0.09 | 0.0187 |
| CD68 | NE | - | - | CD68 | NE | - | - |
| A/L ratio | NE | - | - | A/L ratio | NE | - | - |

HOMA index (IR)
| | Model $R^2 = 0.55$ | | | | Model $R^2 = 0.45$ | | |
| --- | --- | --- | --- | --- | --- | --- | --- |
| | $\beta$ | Partial $R^2$ | $p$ | | $\beta$ | Partial $R^2$ | $p$ |
| Leptin | 0.427 | 0.46 | 0.0069 | Leptin | 0.490 | 0.40 | 0.0041 |
| Adipose cell size | 0.305 | 0.07 | NS | Adipose cell size | 0.236 | 0.03 | NS |
| CD68 | 0.157 | 0.02 | NS | CD68 | -0.152 | 0.02 | NS |

Fasting insulin
| | Model $R^2 = 0.53$ | | | | Model $R^2 = 0.45$ | | |
| --- | --- | --- | --- | --- | --- | --- | --- |
| | $\beta$ | Partial $R^2$ | $p$ | | $\beta$ | Partial $R^2$ | $p$ |
| Leptin | 0.431 | 0.45 | 0.0072 | Leptin | 0.478 | 0.40 | 0.0048 |
| Adipose cell size | 0.293 | 0.06 | NS | Adipose cell size | 0.255 | 0.03 | NS |
| CD68 | 0.147 | 0.02 | NS | CD68 | -0.154 | 0.02 | NS |

Fasting glucose
| | Model $R^2 = 0.26$ | | | | Model $R^2 = 0.13$ | | |
| --- | --- | --- | --- | --- | --- | --- | --- |
| | $\beta$ | Partial $R^2$ | $p$ | | $\beta$ | Partial $R^2$ | $p$ |
| Adipose cell size | 0.233 | 0.23 | NS | Leptin | 0.321 | 0.13 | 0.0117 |
| Leptin | 0.270 | 0.03 | NS | Adipose cell size | NE | - | - |
| CD68 | NE | - | - | CD68 | NE | - | - |

For abbreviations, see legends to Tables 1 and 2; NE: not entered; NS: non-significant.

Despite not being collinear, the strong relationship reported between the plasma A/L ratio and adipose cell size could have masked their independent contribution to IR in our regression models **(Tables 3A and 3B)**. We thus chose to perform partial correlation analyses including HOMA-IR, the plasma A/L ratio, and SCABD or OME adipose cell size. When including OME adipose cell size, the correlation between A/L ratio and HOMA-IR remained significant (partial r=-0.315; p=0.024). In contrast, when including SCABD adipose cell size, the correlation between A/L ratio and HOMA-IR was lost (partial r=-0.253; NS).

However, as plasma A/L ratios and ranges (Table 1) were markedly higher from those of other cohorts of women in which secretory adiposopathy was assessed [26,27,29], the contribution of each adipokine to IR markers was further tested. Plasma leptin levels were positively related to fasting insulinemia and HOMA-IR (0.70<rho<0.73; p<0.0001), and to a lesser extent to fasting glycemia (rho=0.34; p<0.05). Whether these relationships were independent of the associations between circulating L and adipose cell size was also investigated. As these relationships did not reach collinearity (0.56<rho<0.67; p<0.0001), the plasma A/L ratio was replaced by L levels in the regression models where adipocyte size was entered, to verify which of the two variables was the best predictor of insulin resistance markers (**Table 3C**). Leptinemia was the only predictor of HOMA-IR and fasting insulinemia when adipose cell size and CD68 expression in both depots were entered into the model. Circulating L also predicted fasting glycemia, considering adipose cell size and CD68 expression in the OME depot, only.

Finally, as our participant sample showed plasma A/L ratios and ranges well above values already reported in other cohorts of women [26,27,29], we hypothesized that extreme values could be confounding our results. We thus removed those extreme values (5 lowest and 5 highest) from correlation and multiple linear regression analyses. The “corrected” A/L ratio (mean ± SD = 0.99 ± 1.48; median = 0.33) which more resembled to that of other cohorts, correlated to fasting insulinemia and HOMA-IR (−0.58<rho<-0.55; p<0.0005) but to a lesser extent when compared to original values (−0.63<rho<-0.60; p<0.0001). By removing extreme values of the A/L ratio, the latter became the only predictor of fasting insulinemia and HOMA-IR (−0.425<β<-0.416; p=0.01) with adipose cell size and OME CD68 mRNA levels in regression modelling. Using the “corrected” leptinemia values did not change the capacity to predict IR markers (0.39<β<0.68; p≤0.05).

## Discussion

The main objective of this study was to compare the contribution of three markers of AT dysfunction, *i.e*., the plasma A/L ratio, adipose cell size and AT macrophage infiltration, to the variance in cardiometabolic risk factors and more particularly glucose homeostasis in women of varying age and obesity. Taken together, our data show that, in our study population, combined adipose cell hypertrophy at the SCABD or OME level and secretory adiposopathy (a low plasma A/L ratio) lead to a large increase in IR reflected by the HOMA-IR index.

In the present study, only limited and modest relationships were observed between OME CD68 and abdominal AT areas, the lipid-lipoprotein profile or glucose homeostasis. In contrast, SCABD CD68 was a better correlate with these outcomes. However, previous reports have shown positive relationships between total adiposity and body fat distribution and the expression of CD68, CD11b and CD11c in SCABD or in OME AT [16,17]. Our results were not in agreement with the findings of Harman-Boehm et al. who reported in a mixed cohort that the number of CD68+ macrophages infiltrating OME but not SCABD AT was associated with BMI and waist circumference [30]. These differences between our results and others could be explained by cohort composition (men and women vs women, number of participants) or the level of obesity and overall metabolic health, as OME AT macrophage infiltration is particularly exaggerated with centrally distributed obesity [30].

Enlarged fat cell size is considered less metabolically favorable and is generally associated with several of the metabolic abnormalities observed with obesity, including IR [9,31]. For example, men and women with SCABD or OME adipose cell hypertrophy displayed deteriorated insulin sensitivity and increased plasma insulin levels [32]; and women with larger-than-predicted adipocytes in either OME or SCABD AT were characterized by an increased HOMA-IR index [33]. Our data agree with this previous work, as fasting insulin, and glucose levels as well as IR were found to correlate with SCABD and OME adipose cell sizes. Though several studies suggest that OME adipocyte hypertrophy has more deleterious effects on cardiometabolic outcomes when compared to SCABD adipose cell hypertrophy [8,9], the latter has nonetheless been associated with systemic IR in both men and women [31]. In our study, SCABD and OME adipose cell sizes were roughly equivalent predictors and correlates of abdominal AT areas, lipid-lipoprotein profile, and glucose homeostasis. This is perhaps not surprising, given the strong correlation we observed between SCABD and OME adipocyte sizes (rho = 0.78, p <0.0001), in agreement with previous work showing that mean cell size from all anatomical locations generally increases along with adiposity [10].

With regards to the relationship between AT morphology and secretory adiposopathy, we observed a shift toward larger cell sizes with deteriorated secretory function. This was especially true in the SCABD AT. This likely results from the greater impact of varying leptin levels compared to those of adiponectin when calculating the A/L ratio. Indeed, leptin is mainly produced by SCABD AT, and SCABD fat cell size is associated with leptin in both non-diabetic and T2D men and women [34]. As the main source of adiponectin, whose smaller range of values in our sample limits its influence on the A/L ratio when compared to leptin, it is not unexpected that we observed a greater overlap of the distribution curves for OME adipocyte size. Interestingly, the bimodal distribution in our high A/L ratio subgroup implies the presence of both small and large adipocytes in the OME depot of women with a favorable AT secretory profile. This bimodal size distribution was also observed in adipocytes from visceral (*i.e*., omental and mesentery fat depots) of non-diabetic individuals, while it was attenuated in T2D patients with a shift to larger sized cells [35]. We used regression analyses between SCABD or OME fat cell size and the corresponding abdominal fat areas to assess the significance of hyperplasia in AT secretory function. In a complementary analysis, participants were stratified in two subgroups according to the residuals of this regression (either above or under the 95% confidence interval), in which women with a positive residual were considered as having AT hypertrophy, as women with a negative residual were considered as having AT hyperplasia. Our results revealed no difference in the A/L ratio between the two subgroups (data not shown), suggesting that hyperplasia did not impact the association between AT morphology and secretory adiposopathy.

Our data showed that a low plasma A/L ratio was positively associated with several variables of the lipid-lipoprotein profile, as well as fasting insulin levels and HOMA-IR. This adds to a growing number of reports showing associations between secretory adiposopathy and IR markers. For example, we have already shown such relationships with HOMA-IR as well as insulin area under the curve (AUC) in response to an oral glucose tolerance test in healthy women with obesity [26]. Our group further reported differences in fasting insulinemia, insulin AUC, HOMA-IR and HOMA β-cell indices, as well as in the Stumvoll index (which reflects peripheral IS) between postmenopausal women with low and high A/L ratios [36]. Our data showing that women characterized by a low A/L ratio present higher fasting insulinemia and HOMA-IR index reinforce these observations. Furthermore, our data show that women with a low A/L ratio had higher TAG and lower HDL-cholesterol levels as well as a higher cholesterol/HDL-cholesterol ratio, a finding concordant with previous observations of Vega and Grundy [24].

Our group has previously reported that the A/L ratio was an independent predictor of IR and IS in pre- and post-menopausal women characterized by overweight to moderate obesity [26], and an independent predictor of IS in men without T2D [28]. Although the multiple regression models in these studies used waist circumference as an index of adiposity, this measurement cannot distinguish subcutaneous from visceral adiposity [37], or adipocyte morphology. In the present study, while we did observe an independent contribution of the A/L ratio to IR in some multiple regression models, this was lost when SCABD or OME cell size was added to the models. Though this shows that adipocyte cell size is likely a better predictor of IR, conceptually it can be argued that secretory adiposopathy could further add to IR beyond cell morphology. Also, because a strong (but not collinear) correlation was observed between the A/L ratio and adipose cell size, this could have masked independent effects in our models given the limited sample size, a notion supported by our partial correlation analyses data. We thus chose to explore the impact of combined secretory adiposopathy and adipose cell hypertrophy. Results presented in Figure 4 clearly show that the highest HOMA-IR values were observed in women presenting the largest SCABD or OME adipose cell size and the lowest A/L ratio. We also observed stronger correlations between IR markers and circulating L than with the A/L ratio or adipose cell size. Accordingly, leptinemia was a better predictor of IR than both adipose cell size and arguably the plasma A/L ratio, in this cohort. This is in contrast with other studies conducted in women, where the A/L ratio was a better predictor of IR than A or L alone and could be explained by the heterogeneity of our participant sample who were characterized by range and plasma A/L ratios exceeding the one of other studies [26,27,29] probably because of some extreme values due to the large range of adiposity. By removing those extreme (5 lowest and 5 highest) values from our analysis, the “corrected” plasma A/L ratio subsequently predicted some IR markers. Nonetheless, leptin stayed a better predictor of IR markers than the A/L ratio.

Although our study presents many strengths such as the comparison of three distinct markers of AT dysfunction, the determination of body fatness using DEXA, the “gold standard” methodology and of regional fat distribution with computed tomography, as well as the large range of age and total adiposity of our sample, some limitations merit mention.First, as our study included Caucasian pre-, peri and postmenopausal women, results cannot be examined according to the hormonal status (and/or the use of HRT) of our cohort population or extrapolated to men or to different age and ethnic groups. Second, as we measured only the expression of macrophage markers, we cannot assert with certainty that data are equivalent to cell infiltration within AT [16,17]. Third, because AT contains adipocytes and the stromal vascular fraction, heterogeneity in the proportion of each cell type [38] might explain the varying macrophage gene expression pattern observed between ATs, without actual differences occurring within a given cell type between depots. It would be thus interesting to evaluate the proportion of macrophages by immunohistochemistry. Finally, one limitation of our study is the use of collagenase digestion as a single measurement of cell size. This approach underestimates the proportion of very small cells and can generate mean cell sizes that are different from those obtained by other approaches [39]. We suggest that because our analysis is based on collagenase digestion, average cell size should be the predominant variable in our analysis because it is the most accurate parameter derived from this approach.

Among the perspectives, although the plasma A/L ratio is considered a marker of AT secretory dysfunction [23], it remains of interest to assess if specific fat depot A/L mRNA or protein levels would be better predictor of whole-body IR. This could further our understanding of the contribution of different ATs to metabolic health through secretory adiposopathy.

## Conclusion

Our study shows that secretory adiposopathy, assessed as the plasma A/L ratio, is a good predictor of glucose homeostasis, and more specifically IR. Adipose cell hypertrophy, regardless of the fat depot, combined with a low plasma A/L ratio appears to be related to increased IR, in our sample of women of varying age and adiposity. However, leptin was a better predictor of IR markers than both adipose cell size and the A/L ratio in this sample of women. Further studies performed in a larger cohort are clearly warranted to confirm these observations and clarify the potential usefulness of the A/L ratio as a predictor of IR.

## Participants and Methods

### Study Participants

The study included 67 Caucasian women, aged 40-62 years and who underwent abdominal gynecological surgery at the Laval University Medical Center. Women underwent subtotal (n=3) or total abdominal hysterectomies (n=30), some with salpingo-oophorectomy of one (n=11) or two (n=23) ovaries. Reasons for surgery included one or more of the following: menorrhagia/menometrorrhagia (n=33), myoma/fibroids (n=44), incapacitating dysmenorrhea (n=11), pelvic pain (n=3), benign cyst (n=15), endometriosis (n=11), adenomyosis (n=2), pelvic adhesions (n=4), benign cystadenoma (n=1), endometrial hyperplasia (n=5), polyp (n=3) or ovarian thecoma (n=1). Hormonal status was available for 65 women: 2 of 31 premenopausal and 3 of 15 perimenopausal women used hormone replacement therapy (HRT); and only one of 11 postmenopausal women was under HRT for more than 12 months. The status of the other 8 women was uncertain (n = 6) or undetermined (n = 2). This study was approved by the medical ethics committees of Laval University Medical Center and IUCPQ (approval number # 21049). All participants provided written informed consent before their inclusion in the study.

### Body fatness and body fat distribution measurements

Tests were performed on the morning of or within a few days before or after surgery. Measures of total body fat mass, fat percentage and lean body mass, were determined by dual energy X-ray absorptiometry (DEXA), using a Hologic QDR-2000 densitometer and the enhanced array whole-body software V5.73A (Hologic Inc., Bedford, USA). Measurement of abdominal subcutaneous and visceral AT cross-sectional areas was performed by computed tomography, using a GE Light Speed 1.1 CT scanner (General Electric Medical Systems, Milwaukee, USA) and the Light Speed QX/I 1.0 production software, as previously described [40]. Participants were examined in the supine position and both arms stretched above the head. The scan was performed at the L4-L5 vertebrae level with a scout image of the body to establish the precise scanning position. The quantification of visceral AT area was done by delineating the intraabdominal cavity at the internal-most aspect of the abdominal and oblique muscle walls surrounding the cavity and the posterior aspect of the vertebral body using the ImageJ 1.33u software (National Institutes of Health). AT was highlighted and computed using an attenuation range of −190 to −30 Hounsfield units. The coefficients of variation between two measures from the same observer (n = 10) were 0.0, 0.2, and 0.5%, for total, subcutaneous, and visceral AT areas, respectively.

### Adipose tissue sampling and adipocyte isolation

Superficial SCABD and OME AT samples were collected during the surgical procedure (detailed in Participants’ section) and were immediately sent to the laboratory in phosphate buffer saline preheated at 37 °C. SCABD fat samples were collected at the site of a transverse lower abdominal incision, and OME AT was collected from the distal portion of the greater omentum. Adipocyte isolation was performed with a portion of the fresh biopsy and the remaining tissue was immediately frozen in liquid nitrogen and stored at −80 °C for subsequent analyses. AT samples were digested with collagenase type I in Krebs-Ringer-Henseleit (KRH) buffer for 45 min at 37 °C, according to a modified version of the Rodbell method [41,42], as previously described [41,42]. Adipocyte suspensions were filtered through nylon mesh and washed three times with KRBH buffer. The pore size of the mesh used for filtration was 500 μm. Mature adipocyte suspensions were visualized using a contrast microscope attached to a camera and computer interface for cell size measurements. Pictures of cell suspensions were taken, and the Scion Image software was used to measure the size (diameter) of 250 adipocytes for each tissue sample. Average adipocyte diameter was used in analyses.

### Lipid-lipoprotein profile

Blood samples were obtained after a 12-hour fast on the morning of surgery. Cholesterol and triacylglycerol (TAG) levels were measured in plasma and lipoprotein fractions with a Technicon RA-analyzer (Bayer, Etobicoke, Canada) using enzymatic methods, as described previously [43]. The high-density lipoprotein (HDL) fraction was obtained by precipitation of low-density lipoproteins (LDL) from the infranatant with heparin and MnCl_2_ [44]. The cholesterol content of the infranatant measured before and after precipitation, and the concentration of LDL-cholesterol was obtained by difference.

### Glucose homeostasis and adipokines

Fasting glucose and insulin levels were measured in blood samples obtained on the morning of surgery. Plasma glucose was measured using the glucose oxidase method and plasma insulin by radioimmunoassay (Linco Research). The HOMA insulin resistance (HOMA-IR) index was calculated using fasting insulin (μU/mL) x fasting glucose (mmol/L)/22.5 [45]. Plasma leptin (Linco Research) and adiponectin (B-Bridge International) levels were measured by enzyme-linked immunosorbent ELISA assays from pre-surgery blood samples.

### mRNA determination by quantitative real-time reverse transcriptase polymerase chain reaction

Total RNA was isolated from SCABD and OME AT using the RNeasy lipid tissue extraction kit and on-column DNase treatment (Qiagen) following the manufacturer’s recommendations. RNA quality and concentration were assessed using the Agilent Technologies 2100 Bioanalyzer (Agilent). Complementary DNA was generated from 4 ug of total RNA with 50 ng of random hexamers, 300 ng of oligo dT18, and 200 U of Superscript II Rnase H-RT (Invitrogen Life Technologies) and purified with QIAquick PCR Purification Kit (Qiagen). Real-time complementary DNA amplification was performed in duplicate using the Light Cycler 480 (Roche Diagnostics, Indianapolis, USA) and SYBR Green I Master (Applied Biosystems, Foster City, USA), as follows: 95°C for 10 seconds, 60°C to 62°C for 10 seconds, 72°C for 14 seconds, and then 76°C for 5 seconds (reading) repeated 45 times. Target gene amplifications were normalized using expression levels of ATP synthase O subunit (ATP5O) as the housekeeping gene. Expression levels of this gene were not associated with age and adiposity in our study sample. Similar results were obtained using other housekeeping genes. Primer sequences for CD68 (NM_001251; sense: 5′-GCAGCAACTCGAGCATCATTCTT-3′; anti-sense: 5′-CGAGGAGGCCAAGAAGGATCA-3′), CD11b (NM_000632; sense: 5′-TTCCAGAACAACCCTAACCCAAGATC-3′; anti-sense: 5′-ATCGCCAAACTTTTCTCCATCCG-3′), and CD11c (NM_000887; sense: 5′-GGCCATGCACAGATACCAGGT-3′; antisense: 5′-CTGGGGGTGCGATTTTCTCTG-3′), were designed using Gene Tools (Biotools), as previously described [45].

### Statistical analysis

Statistical analyses were performed using JMP software (SAS Institute, Carry, NC, USA). Data were considered statistically different when p < 0.05. Non-normally distributed variables were log10 transformed for parametric analyses. Spearman correlations were used to assess relationships between variables. The Kolmogorov-Smirnov test was used to investigate differences in adipocyte size distributions between patients with low vs high plasma A/L ratio tertiles. Partial correlations were used to investigate relationships between the plasma A/L ratio, SCABD or OME adipocyte size, and HOMA-IR. Finally, the A/L ratio and SCABD or OME adipose cell size were divided into tertiles (high, mid, and low), and 3-D graphs were used to highlight the impact of combined secretory adiposopathy and fat cell size on HOMA-IR. Stepwise (mixed) regression modeling was used to assess the independent contributions of the retained variables to IR (HOMA index), and to fasting insulin and glucose levels. Two AT dysfunction markers, the plasma A/L ratio and SCABD or OME CD68 mRNA levels, and one lipid-lipoprotein marker, TAG levels, were chosen based on the strength of simple correlations with the predicted variables and included in our first model. Plasma TAG levels were replaced by the SCABD or OME adipose cell size in a second model. Due to the significant relationships between plasma L levels and glucose homeostasis, the A/L ratio was replaced by circulating L in a third model.

## Declarations and ethics statements

### Ethical Approval

This study was approved by the medical ethics committees of Laval University Medical Center and IUCPQ (approval number # 21049).

### Informed consent from participants

All participants provided written informed consent before their inclusion in the study.

### Data availability statement

All relevant data are within the paper and its Supporting Information files.

### Author contributions

EJT performed the statistical analyses, interpreted the results, and developed the draft of the manuscript. NC helped EJT in conducting analyses and interpreting results. PM and DRJ supervised the study and provided a scientific framework for the study. MP participated to data collection, data base organization and ethics approval. AT supervised the clinical study in collaboration with the medical team. All authors read and approved the final manuscript.

## Acknowledgements

The cooperation of study participants involved in this study is greatly appreciated.

